# Causal Inference Engine: A platform for directional gene set enrichment analysis and inference of active transcriptional regulators

**DOI:** 10.1101/698852

**Authors:** Saman Farahmand, Corey O’Connor, Jill A. Macoska, Kourosh Zarringhalam

## Abstract

Inference of active regulatory mechanisms underlying specific molecular and environmental perturbations is essential for understanding cellular response. The success of inference algorithms relies on the quality and coverage of the underlying network of regulator-gene interactions. Several commercial platforms provide large and manually-curated regulatory networks and functionality to perform inference on these networks. Adaptation of such platforms for open-source academic applications has been hindered by the lack of availability of accurate, high-coverage networks of regulatory interactions and integration of efficient causal inference algorithms. In this work, we present CIE, an integrated platform for causal inference of active regulatory mechanisms form differential gene expression data. Using a regularized Gaussian Graphical Model, we construct a transcriptional regulatory network by integrating publicly available ChIP-Seq experiments with gene-expression data from tissue-specific RNA-Seq experiments. Our GGM approach identifies high confidence TF-gene interactions and annotates the interactions with information on mode of regulation (activation vs. repression). Benchmarks against manually-curated databases of TF-gene interactions show that our method can accurately detect mode of regulation. We demonstrate the ability of our platform to identify active transcriptional regulators by using controlled *in vitro* overexpression and stem-cell differentiation studies and utilize our method to investigate transcriptional mechanisms of fibroblast phenotypic plasticity.

## INTRODUCTION

Technological advancements in high-throughput sequencing have made it possible to measure expression of genes at relatively low cost. However, the direct measurement of regulatory mechanisms, such as transcription factor (TF) activity, in a high-throughput manner is still not readily available. Consequently, there is a need for computational approaches that can identify active regulatory mechanisms from observable gene expression data. The scientific community has developed a multitude of algorithms and biophysical models to study the impact of TF activity on gene expression. Some of these algorithms attempt to infer TF activity and dynamics directly from gene expression data (1, 2). Others rely on biophysical approaches to model expression of genes based on known TF-gene interactions (3). Another class of algorithms, which are the main focus of this work, use prior biological knowledge on biomolecular interactions to link a differential gene expression (DGE) profile to upstream regulators (e.g., TFs) (4, 5, 6, 7). The essential ingredients of these algorithms are (i) a DGE profile, (ii) a network of biomolecular interactions, and (iii) an inference algorithm to query the network.

The DGE profile as obtained from RNA-Seq or microarrays studies is the observable input and quantifies the difference in transcript abundance between two conditions (e.g., healthy vs. disease, stimulated vs. not stimulated, etc.). The network of biomolecular interactions encapsulates the prior biological knowledge. The accuracy and ability of inference algorithms to identify upstream molecular drivers of observed DGE profiles rely to a large degree on the quality and coverage of the network and availability of auxiliary information on interactions within the network. There are several sources of publicly available protein-protein interactions and signaling pathways (e.g., STRINGdb (8), Pathway Commons (9), Kyto Encyclopedia of Genes and Genomes (KEGG) Pathways (10), etc.). In case of regulatory networks, Ingenuity (www.ingenuity.com) provides a high-coverage manually-curated network of regulatory interactions in an integrated platform, *Ingenuity Pathway Analysis tool* (4). Among other things, this platform provides pathway inference and enrichment analysis functionality and network visualization tools. However, the Ingenuity tools are inaccessible for the majority of academic applications and there is a need for freely available alternatives for academic purposes.

Several approaches for reconstruction of regulatory networks form gene expression data have been proposed by the scientific community (11, 12). Public sources of gene regulatory interactions are either computationally derived, based on biological experiments, or manually curated from biomedical literature (13, 14, 15). These sources include the Transcriptional Regulatory Element Database (TRED) (16), the Transcription Regulatory Regions Database (TRRD) (17), and Transcriptional Regulatory Relationships Unraveled by Sentence-based Text Mining (TRRUST) (18). These databases provide valuable information on gene regulatory mechanisms, but drawbacks exist. The scope of the databases containing experimentally validated interactions are very small, and cover only a fraction of TF-gene interactions. On the other hand, databases of computationally predicted and expression-driven interactions are typically very noisy. Importantly, the majority of the databases do not report the direction of regulation (activation vs. repression) - which is crucial to understanding the functional behavior of the cell.

In this work, we present Casual Inference Engine (CIE), a platform for active regulator inference on biological networks consisting of a web-server and a user-friendly `R`-package. The platform provides various inference models, including methods based on Fisher’s exact test (enrichment test) as well as directional enrichment models that can utilize information on mode of regulation (6, 7). Moreover, we present an approach based on regularized Gaussian Graphical Models (GGM) to construct an accurate and high-coverage annotated networks of TF-gene regulatory interactions. We achieve this by integrating publicly available high-throughput ChIP-Seq experiments deposited in ChIP-Atlas (19) with tissue specific gene expression data from GTEx (20). We show the consistency and accuracy of our reconstructed network by benchmarking against manually curated interactions in the TRRUST database. We demonstrate the utility of our platform in identifying active regulators using controlled *in vitro* overexpression studies as well as more complex gene expression data from a stem cell differentiation experiment. Additionally, we show how our platform can assist in identifying novel transcriptional regulatory mechanisms using gene expression data from primary prostate fibroblast cells stimulated with TGF*β* and CXCL12. Although our focus in this work is on transcriptional regulatory networks, the `R`-package provides functionality to perform inference on any type of user provided network. The CIE platform provides higher-order pathway enrichment analysis on identified active regulators using Reactome pathways (21). Figure 1 shows a schematic overview of the CIE platform.

**Figure 1.**
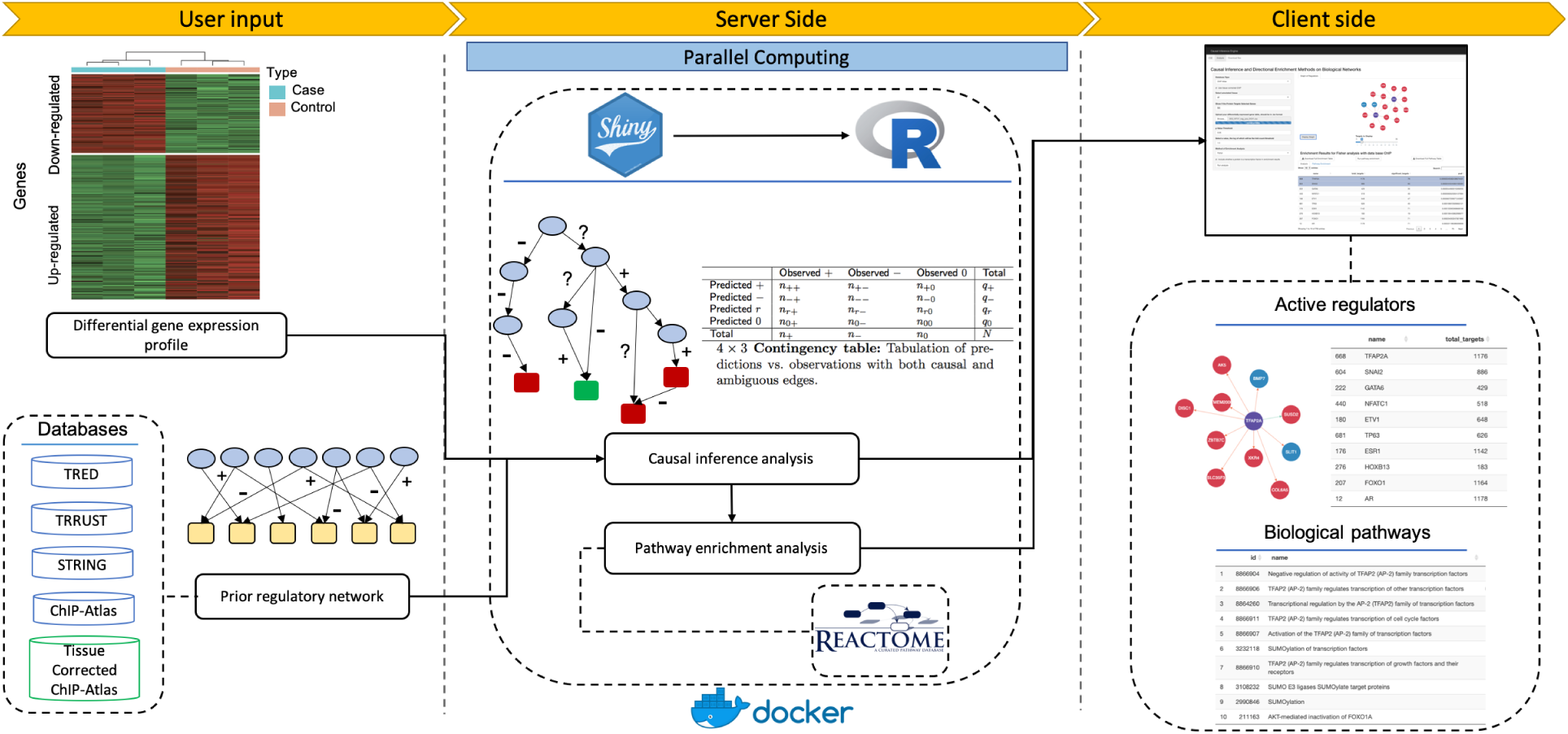
CIE platform for inference of active regulators. CIE takes a user provided DGE profile as input (left panel). User selects a prior regulatory network from the provided databases. In the server side (middle panel), the input is processed and the Shiny web-server starts `R` scripts to perform the inference analysis based on the user selected options. Predicted active regulators associated with the DEG profile are displayed with interactive graphics and are downloadable in table format (right panel). CIE also offers pathway enrichment analysis by mapping the inferred regulators to pathways from the Reactome database.

## MATERIALS AND METHODS

### Regulatory network

The interaction network can be viewed as a graph *G* = (*V,E*), consisting of a set of nodes *V* (biological entities) and a set of edges *E* =(*u,v*). The network is not limited to interactions between TFs and genes and can include other types of interactions (e.g., interactions between compounds and proteins, protein-protein interactions, etc.). Some of these edges may be associations (an undirected edge *u* − *v*, indicating change in *u* is correlated with change in *v*), causal (a directed edge *u* → *v*, indicating that *u* regulates *v*) or signed causal (with + or −, indicating mode of regulation).

### Databases

We utilized three sources of TF-gene interaction networks. Our criterion for inclusion was that they either must include high-confidence, manually curated interactions with literature support or must have direct experimental evidence. These sources are:

(A) The TRRUST database: TRUSST (18) is a manually-curated database of human transcriptional regulatory network derived from PubMed articles with partial information on mode of regulation. It contains 9,396 regulatory interactions between 795 human TFs and 2,067 target genes.

(B) The TRED database: TRED (16) is an integrated repository for both *cis-* and *trans-* regulatory elements in mammals with experimental evidence. It includes a total of 6,726 interactions between 36 TF and 2,910 genes. These interactions are not annotated with mode of regulation.

(C) ChIP-Seq derived TF-gene interactions: We obtained all publicly available ChIP-Seq data (> 96,000 experiments) that are processed and deposited into ChIP-Atlas (19). A TF-gene interaction network was assembled by merging all experiments and applying various filters for peak signal intensity (0-1,000) and distance to the transcription start site (TSS) (1kb, 5kb, and 10KB). These filters are integrated in the CIE platform and can be applied interactively in the web-app. For example, peak intensity score of 500 and distance of 5kb to TSS results in 185,271 interactions between 642 TFs and 16,148 target genes. Note that ChIP-Seq network does not directly provide information on mode of regulation and all TF-gene interactions in this database are unannotated.

(D) STRINGdb: In addition to TF-gene interactions, we also included Protein-gene interactions from STRINGdb (8). STRINGdb includes protein-protein interactions (PPI) from various sources including curated, experimentally supported, and computationally-derived interactions. Some of the interactions are causal and annotated, but most interactions in the STRINGdb are undirected. For each undirected PPI *u*−*v*, we constructed two directed interactions *u* → *v* and *v* → *u*.

### Differential Gene Expression Profiles

We used several DGE profiles from microarray and RNA-Seq experiments to evaluate the utility of our platform. For microarray data, gene expression profiles were normalized and differentially expressed genes were computed using the `R` limma package (**?**). We applied a 1.3 absolute value fold change and *<* 0.05 FDR corrected p-value filter for selecting differentially expressed genes. RNA-Seq data was processed using the HISAT2 (22) pipeline and differentially expressed genes were identified for each treatment the edgeR package (23). Similar filters for FDR and fold change were applied to identify differentially expressed genes. The data sets that we utilized for our evaluation are:

#### Controlled overexpression experiment

We utilized three datasets from (24), in which recombinant antiviruses were used to infect normal human epithelial cell in order to overexpress specific oncogenes. The over expressed genes are E2F3, c-Myc and H-Ras. There are 272, 220 and 268 differentially expressed genes compared to the WT in the experiments respectively.

#### Stem cell directed differentiation

We used a time-course in vitro differentiation model of pancreatic beta cell development from (25). NEUROG3+ Pancreatic progenitor cells convert to NKX2-2+ endocrine cells, which are able to further differentiate into fully-functional insulin producing cells upon implantation into mice (26). A total of 1,000 differentially expressed genes were identified from this dataset.

#### fibroblast phenotypic plasticit

We utilized data from RNA-Seq experiments performed on prostatic stromal fibroblasts stimulated with Vehicle, TGF*β*, and CXCL12 (27). A total of 10,032 differentially expressed genes were identified in fibroblasts treated with TGF*β* or CXCL12. The DGE profile of TGF*β* and CXCL12 were 75% similar (7,502 transcripts). A total of 1,012 (10%) were induced by TGF*β* treatment only and 1,357 (13%) by CXCL12 treatment only. 161 (2%) were differentially regulated in opposite directions by CXCL12 and TGF*β*.

### Construction of transcriptional regulatory networks

The network constructed from ChIP-data is noisy as the experiments are performed under various conditions in different cell lines. Moreover, the interactions are not annotated with mode of regulation (activation vs. repression). To reduce the noise and annotate the interactions, we utilized a regularized Gaussian Graphical Model (glasso) (28) to integrate the ChIP-derived network with tissue specific RNA-Seq data obtained form the Genotype-Tissue Expression project (The GTEx Consortium) (20). Processed normalized gene expression values were obtained from (29), where GTEx RNA sequencing reads from 15 tissues from 2,585 paired-end RNA-Seq samples were re-processed, uniformly realigned, and normalized to remove batch effects and tissues with low number of samples. Every ChIP-derived interaction was taken account without any filters (≈4*×* 10^6^ interactions). To construct the tissue-specific annotated regulatory networks, we estimated a sparse covariance matrix using each tissue-expression data separately, while softly enforcing ChIP-derived interaction using an *l*_1_ penalty matrix. Gene expression was log transformed prior to analysis. Only protein-coding genes were utilized and genes with no on-to-one map between Ensemble ID and HGNC symbol were excluded.

The process of constructing a posterior network form gene expression data and ChIP-network is as follows. Let *S* denote the empirical covariance matrix estimated from the RNA-Seq expression data for a given tissue, Σ be the (unknown) covariance matrix, and Θ= Σ^−1^ be the precision matrix. Glasso directly estimates the precision matrix Θ by maximizing the *l*_1_ penalized log likelihood

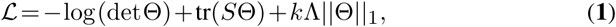

on the space of positive semi-definite matrices. Here Λ is a shrinkage parameter matrix and *k* is a scaler tuning parameter. The prior ChIP-network structure was incorporated in the penalty as follows. First, we constructed an adjacency matrix *A* from the ChIP-network. The rows and columns of *A* were arranged by TFs first and then by genes. If there is a connection between transcription factor *i* and gene *j* the corresponding entry in the adjacency matrix is set to 1 (i.e., *A*_*ij*_ = 1), and otherwise it is set to 0. The penalty matrix Λ has the same size as the adjacency matrix. The entries of this matrix are constant values and are set to differentially penalize the connections based on the information in the adjacency matrix as follows:

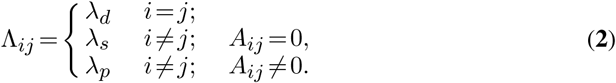

In our implementation, diagonal elements were not penalized (i.e., *λ*_*d*_ = 0). Interacting TFs and genes (i.e., ., *A*_*ij*_ = 1) were penalized by a small nonzero value (*λ*_*s*_ = 0.05) and non-interacting pairs were penalized by a relatively large value (*λ*_*p*_ = 0.5). The matrix was then scaled by a constant value *k*. We utilized a path of values ranging from 1 to 6 with step size 0.1 for *k*. For each scaling value, we fitted the model by maximizing the log likelihood and calculated the corresponding precision matrix Θ, from which a posterior regulatory network was constructed based on the conditional independence property of GGMs (30). More precisely there is a connection between TF *i* and gene *j* if and only if Θ_*ij*_ ≥ *ϵ*. The threshold value *ϵ* was selected 1 − *e* 4 empirically as a small value. For each posterior network, we calculated the scale-free property using the R-squared (*R*^2^) value between log(*p*(*d*)) and log(*d*), where *p*(*d*) represents the proportion of nodes in the network with *d* interactions (31, 32). We chose a value of *k* for each tissue that generated the highest *R*^2^ value. Figure 2 illustrates the approach. Once the final posterior network was constructed, the signs of the interactions were determined using the partial correlation matrix:

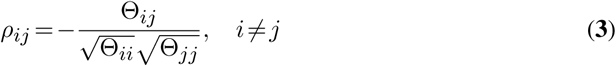

**Figure 2.**
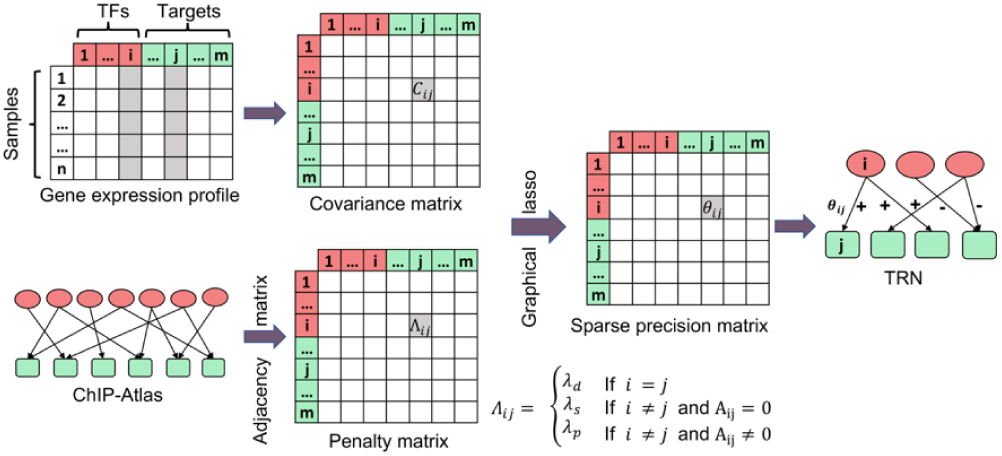
Assembly of tissue-specific regulatory networks. The empirical covariance matrix was estimated from GTEx tissue-specific gene expression data. ChIP-Atlas prior network was converted into an adjacency matrix of prior TF-gene interactions and softly enforced into graphical lasso using a penalty matrix Λ with various degrees of stringency. The sparse precision matrix Θ was estimated by optimizing the penalized log likelihood, from which the posterior network was constructed. The precision matrix encodes the direct interaction between entities in the network. A non-zero value Θ_*ij*_ ≥ *ϵ* indicates that there is an interaction between *i*th TF and *j*th target gene. The sign of interaction (activation vs. repression) is calculated from the partial correlation matrix.

Note that the connections in each posterior network are supported by both ChIP-Seq data as well as by partial correlation of gene expression values. A total of 15 tissue-specific posterior network were constructed using this process. Additionally, we examined the overlap between these networks to identify the connections that appear across multiple tissues. Such connections can be viewed as consistent universal interactions. We refer to these network as merged networks. In our implementation, we constructed merged network using interactions that are consistent across 2, 3, 4, or 5 tissues.

### Inferring active regulators

CIE provided several (directional and un-directional) enrichment tests to query the networks and identify transcriptional regulators from a user provided DGE profile. The starting point of the inference is selection of one of the causal networks of interactions that are provided by CIE. The type of inference depends on the availability of information on the mode of regulation in the causal graph. The first method is the Fisher’s exact test or the enrichment scoring (ES) statistic, which is the standard for gene set enrichment analysis (33). This method does not take information on mode regulation into account. The next enrichment method is Ternary scoring (TS) statistic proposed by Chindelevitch et. al. (7). This method is suitable for fully annotated networks. For networks with a mixture of annotated and unannotated edges, we utilized the Quaternary scoring (QS) statistic proposed by Fakhry et. al.(6).

All these methods are build on a common core, calculating the goodness of the fit of the score, which measure the agreement between predictions made by regulators in the graph and the observed DEG profile. For each regulator in the selected network, a contingency table is constructed that tabulates the agreement between the predictions made by the regulator according to the graph and the observations based on the DGE profile. The rows of the table represent the prediction made by the regulator and the columns represent observed differential expression profile. In case of the Enrichment scoring statistic, a 2 × 2 contingency is constructed, with rows representing genes that are predicted to be regulated, or unregulated by the regulator according to the network, and the columns representing genes that are observed to be regulated, or unregulated according to the observed DGE profile. In case of Ternary scoring statistics a 3 × 3 contingency table is constructed, with rows representing genes that are predicted to be upregulated, downregulated, or unregulated by the regulator and the columns represent genes that are observed to be upregulated, downregulated, or unregulated according to the DGE profile. Finally, in case of Quaternary scoring statistic a 4 × 3 contingency table is constructed in a similar manner with an additional row representing regulated or unregulated corresponding to unsigned edges in the network. The statistical significance of the score is calculated using a permutation (generalized hyper-geometric) test and the p-values are reported. Details of these algorithms can be found in the original publications (6, 7).

### The CIE web server

The CIE web server was implemented using `R` Shiny, Docker, ShinyProxy, and a NGINX web server. The applet RCytoscapeJs was used for network visualization. The core of functionality of the web server is driven by the CIE `R` package. The open-source version of Shiny is a single-threaded web server for web applications implementing with `R`. If it is used to run a web application that takes a few seconds at most to load, this will not cause any noticeable impairment. However, CIE takes up to several minutes to load and the open-source Shiny web server cannot allow another user to connect during this time as it can only do one task at a time. To overcome this limitation, we used ShinyProxy and Docker container to allow creation of multiple instances. ShinyProxy detects a new user’s request and starts a Docker container of CIE application specifically for the user. NGINX takes the request from the user and forwards it through reverse proxy to ShinyProxy. The CIE web server is located at https://markov.math.umb.edu/app/cie, and is accessible by all major browsers. The inference result table consists of the regulator’s symbol, total number of target genes, number of the significant target genes (i.e., differentially expressed genes), and the corresponding p-values. The result table can be downloaded in a text format. Users can also run higher-order pathway enrichment analysis by mapping the inferred regulators to Reactome pathways (21).

### The CIE R-package

We also provide CIE `R`-package for offline and local usage available to download at https://github.com/umbibio/CIE-R-Package, under the GNU public license. The package is capable of producing the same plots and results as the web server with a simple functional call and allows for more fine-tune control, automation, and customized input networks. The CIE `R` package is parallelized to provide efficient and fast inference. It utilizes the multidplyr and dopar packages to implement this parallel computation. Multidplyr allows the table of statistics from which enrichment is calculated to be produced quickly, and dopar calculates the p-values in parallel by wrapping a function call to their calculator. Our package is documented and includes a comprehensive manual and instructional vignettes.

## RESULTS

We performed our GGM approach to integrate ChIP-derived network from ChIP-Atlas (19) with tissue-specific gene expression data from GTEx (20). For each tissue, a grid of regularization parameters was applied, and the best network based on the highest R-squared value was selected (SI Table 1).

**Table 1.** Top 20 predicted regulators by CIE on tissue corrected ChIP-network.

| E2F3 |  |  |  | c-Myc |  |  |  | H-Ras |  |  |  |
| --- | --- | --- | --- | --- | --- | --- | --- | --- | --- | --- | --- |
| Name | ES | TS | Reg. | Name | ES | TS | Reg | Name | ES | TS | Reg. |
| E2F1 | 3.9e-8 | 4.9e-11 | up | TP53 | 8.8e-18 | 2.5e-20 | up | FOSL1 | 2.6e-28 | 2.0e-42 | up |
| PSIP1 | 1.7e-6 | 1.6e-6 | up | SSRP1 | 3.5e-14 | 2.2e-22 | up | TP63 | 4.5e-13 | 9.6e-2 | up |
| FOXM1 | 3.7e-6 | 1.5e-8 | up | WDR5 | 1.9e-12 | 5.6e-20 | up | KDM6B | 2.4e-12 | 6.2e-19 | up |
| EZH2 | 3.7e-6 | 2.2e-4 | up | SMARCA4 | 1.0e-11 | 2.7e-12 | up | ELF3 | 2.1e-11 | 1.6e-12 | up |
| BRCA1 | 1.5e-5 | 3.1e-3 | up | E2F4 | 2.1e-9 | 3.3e-17 | up | EGR1 | 3.1e-11 | 3.0e-6 | up |
| E2F2 | 4.3e-5 | 1.1e-5 | up | MYC | 2.1e-8 | 1.9e-5 | up | MAFF | 3.2e-10 | 3.3e-15 | up |
| MYBL2 | 5.7e-5 | 5.6e-4 | up | TRIM28 | 1.7e-7 | 9.6e-14 | up | FOS | 1.7e-9 | 1.0e-5 | up |
| E2F7 | 6.8e-5 | 5.4e-6 | up | TEAD4 | 1.9e-7 | 3.7e-12 | up | JUNB | 2.9e-9 | 1.9e-12 | up |
| ETS1 | 1.2e-4 | 6.6e-3 | up | NFKB1 | 6.2e-7 | 1.7e-6 | up | BCL3 | 6.1e-8 | 2.7e-8 | up |
| CEBPD | 1.0e-3 | 1.8e-2 | down | ILF3 | 8.4e-7 | 3.4e-12 | up | SMAD3 | 6.7e-8 | 7.7e-9 | up |

Columns: Predicted regulator (Name), FDR corrected p-value (Enrichment Score: ES and Ternary Score: TS), predicted direction of regulation (Reg.)

### Summary statistics

We examined the overlap between recovered interactions in each tissue. Figure 3a shows the number of recovered regulatory interactions shared between tissues. Interactions that appear across several tissues are called consistent and reflect non-tissue specific, universal TF-gene interactions. As expected, we observed a significant drop in number of consistent interactions as number of tissues increase. There are a total number of 95,745 interactions between 739 TFs and 14,660 genes that appear across at least 5 tissues. We further examined the consistency of recovered signs (mode of regulation) across tissues. This analysis was carried out on interactions that appear in at least 5 tissues. For each interaction, the proportion of times that the interaction was annotated as positive (or negative) across the tissues in which the interaction appeared was calculated. Completely consistent interactions will have either positive proportion of 1 (i.e., recovered always as positive) or 0 (i.e., recovered always as negative). Figure 3b shows the frequency of the positive proportions. As can be observed the distribution is bimodal, with most interactions recovered consistently as either positive or negative, indicating that TF-gene interactions tend to be activation or repression independent of the tissue. For cross-tissue merged networks, a majority voting scheme was used to annotate inconsistent interactions.

**Figure 3.**
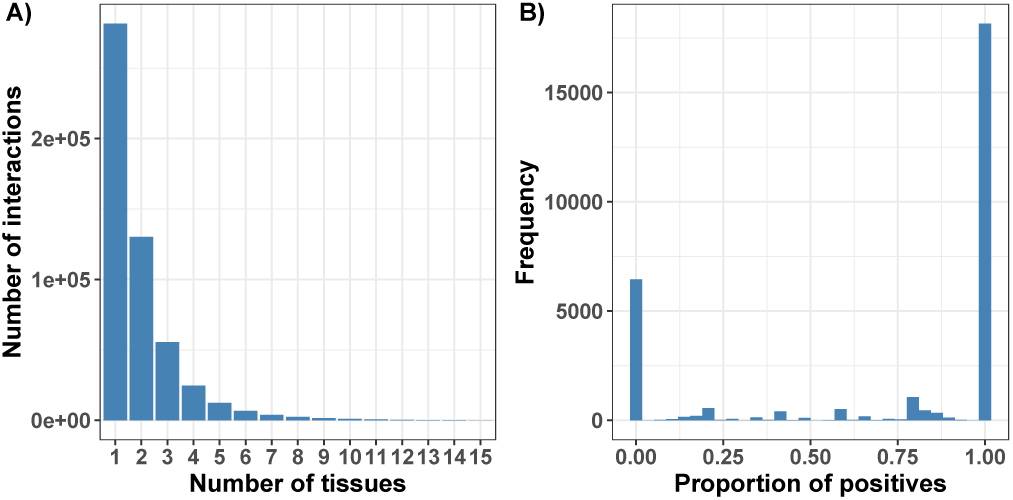
**A)** Number of interactions shared across tissues. **B)** P of positive interactions shared in at least 5 tissues. The bimodal d shows that interactions are consistently annotated as positive (1.0) or negative (0).

### Benchmark results

We compared our inferred mode of regulation with the signs reported in the TRRUST database, a high-quality, manually-curated database of human TF-gene interactions, which can be considered as the gold standard for our purpose. For this benchmark, we merged all annotated interactions across all tissues. Figure 4a shows the number of posterior interactions and their associated sign distribution recovered by the GGM approach. Of the interactions in the ChIP-network, 47% were supported by a tissue specific gene expression data and annotated by our GGM approach (Figure 4a top). The final sign of recovered interactions were decided according to a majority voting scheme resulting in 55% positive and 45% negative interactions (Figure 4a top). We compared the sign of annotated interactions with the signs reported in the TRRUST database. The overlap between the ChIP-network and the TRRUST data base is 3,812 TF-gene interactions (Figure 4a bottom), of which 1,247 are annotated in both TRRUST and the tissue corrected ChIP-network. Further restricting the interactions in the ChIP-network to consistent interactions that appear across at least 5 tissues, results in 147 overlap with annotated TRRUST interactions (see also SI Figures 1 and 2). Figure 4b shows classification performance of these 1,247 and 147 overlapping interactions between tissue corrected and cross 5-tissue merged ChIP-networks with TRRUST. As can be seen the agreement is high in both cases (F1-scores 0.75 and 1.0), demonstrating that our approach is highly accurate in identifying sings of regulation.

**Figure 4.**
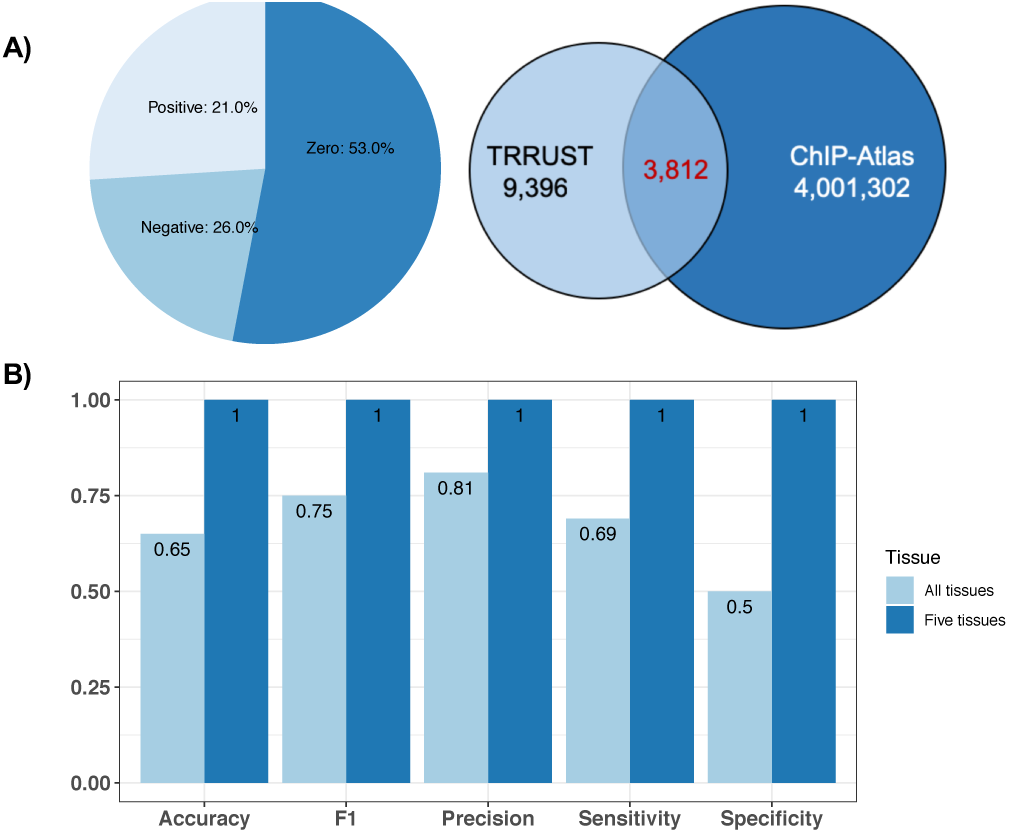
**A)** Summary statistics of annotated interactions in the ChIP-network and overlap with TRRUST database. **B)** Classification performance of annotated interactions in ChIP-network compared with TRRUST.

### Recovering known perturbations in controlled overexpression experiments

To test the performance of our tissue-corrected ChIP-network and inference algorithms, we used the CIE platform to identify drivers of differential expressed genes in controlled overexpression studies. For this analysis, we utilized three differential expression profiles, all of which were obtained by over expressing an oncogene. The genes are E2F3, c-Myc, and H-Ras. The number of differentially expressed genes in experiment are 272, 220 and 268 respectively. Table 1 outlines the top 10 regulators predicted by CIE on each experiment sorted by the FDR corrected p-values of the enrichment statistics (ES). The FDR corrected p-values of the Ternary score (TS) along with the predicted direction of regulation by the Ternary method are also presented in the table.

In the case of E2F3 experiment, E2F1 is returned as the top putative regulator along with E2F2 as another top regulator. E2F1, E2F2, and E2F3 are close related family of transcription factors with very similar roles that function to control the cell cycle and are implicated in cancer (34). The direction of regulation for these factors are correctly predicted as upregulated by Ternary method. Another predicted regulator is EZH2, which is a downstream of the pRB-E2F pathway and is essential for prolifiration and amplified in several primary tumors (35). Interestingly, the Ternary method predicts the direction of regulation of CEBPD as down. It is documented that CEBPD reverses E2F1-mediated gene repression and increased level of CEBPD attenuates E2F1-induced cancer cell proliferation (36). In the c-Myc experiment, the algorithm recovered MYC as one of the top putative regulators and it is predicted to be upregulated. CIE also predicted WDR5, a required interactor of MYC that associates with the same target genes in vivo and is implicated in driving tumorigenesis (37). Finally, in the H-Ras experiment, EGR1 is the top putative regulator returned by the algorithm with a significant p-value and predicted to be upregulated. EGR1 is a key regulator of oncogenic processes and is downstream of H-RAS (38). In all cases the biology behind the recovered regulators is sufficiently evident, demonstrating the ability of the CIE platform and the tissue corrected ChIP-network in recovering correct regulators of differential gene expression.

Note that Ternary (and Quaternary) scores are generally more stringent than the enrichment score. Unlike the enrichment score, they also match the direction of regulation between the network and the DEG profile. Additionally, the Ternary and Quaternary are able to make inference on the direction of perturbation of the predicted active regulators.

### Drivers of stem cell directed differentiation

To test the utility of the CIE platform and the higher-order pathway enrichment on inferred regulators in generating biological insight, we utilized a more complex data set of stem cell directed differentiation of pancreatic beta cells (25) and the mixed signed STRINGdb network with Quaternary scoring method. Figures 5A and B show the CIE causal regulatory inference results. Among the top predictors, Gastrin (GAST), IL6 and NEUROG3 are predicted to be up-regulated by CIE, all of which are involved in the development of the pancreatic endocrine cell lineage (39, 40). Users can interactively select top regulators predicted by CIE and perform pathway enrichment analysis using the Reactome (21) pathways. Figure 5C shows the enriched Reactome pathways returned by CIE using the top 5 predicted regulators. Several significant pathways were acquired through this analysis such as regulation of gene expression in late stage (branching morphogenesis) pancreatic bud precursor cells, which is essentially pointing to the endocrine differentiation of the epithelial cells (41). Furthermore, Regulation of beta-cell development is also identified, which provides a direct link to the transient cellular stages leading to the generation of all pancreatic endocrine cells including insulin-producing beta cells (41). Finally, transcriptional regulation of pluripotent stem cells is identified, which encodes regulatory networks underlying Embryonic stem cells (ESCs) differentiation into any cell type or tissue type in body (42).

**Figure 5.**
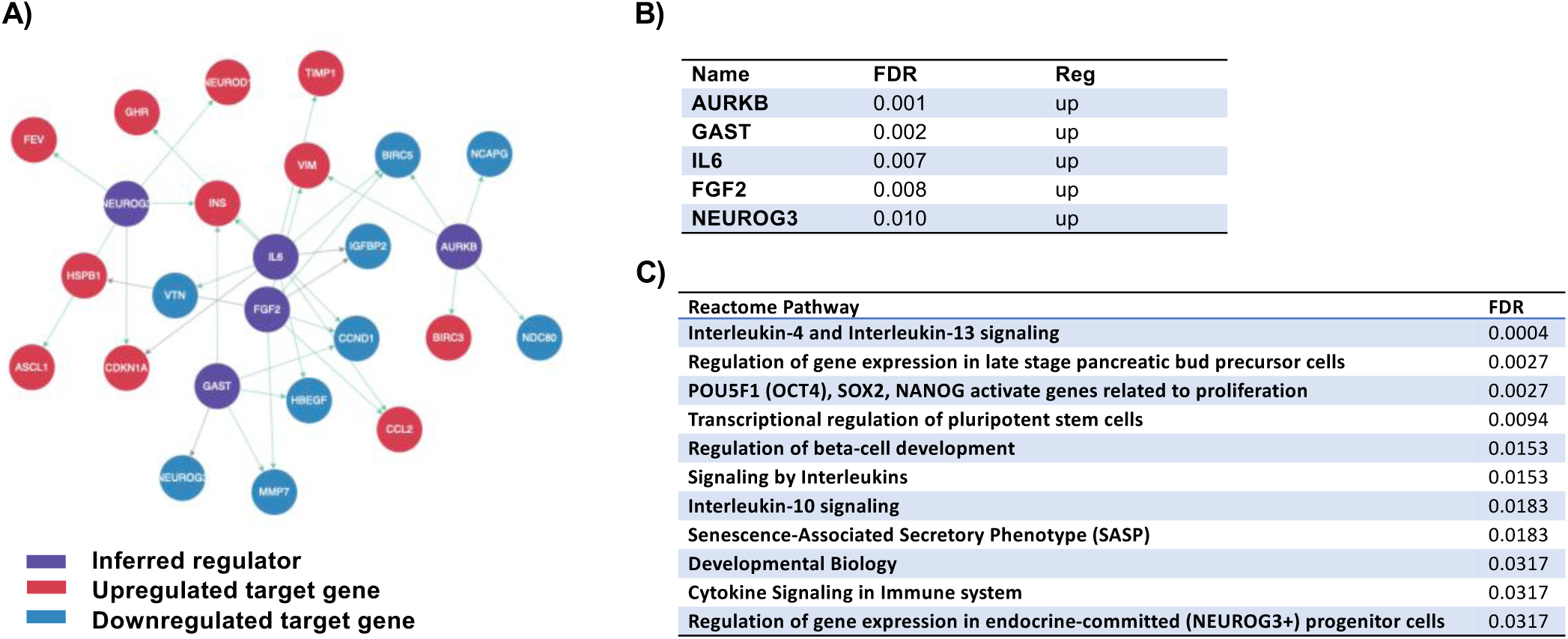
CIE output on differentiation of pancreatic beta cell. **A)** Inferred regulatory network corresponding to top 5 predicted regulators and their target genes. **B)** Table of top 5 predicted active regulators. **C)** Reactome pathways corresponding to the top 5 predicted regulators by CIE.

### Signaling mechanisms underlying fibroblast-to-myofibroblast phenoconversion

In this section we show how the CIE platform can be utilized to test a specific biological hypothesis regarding transcriptional regulators. We demonstrate this use case in the context of fibroblast phenotypic plasticity. Fibroblasts are an abundant cell type within the human connective tissue and play a primary role in secretion of the components of the extracellular matrix (ECM) (43). Fibroblasts have striking similarities with mesenchymal stem cells (MSCs) and share some functions with MSCs, including phenotypic plasticity governed at the genetic level (44). It is known that Fibroblast can phenotypically convert to myofibroblasts in response to pro-fibrotic proteins such as TGF*β* and CXC-type chemokines, secreted by the aging and/or inflammatory cells (27, 45, 46). Phenotypic conversion can occur through two independent cellular signaling mechanisms: One that depends upon TGF*β*/TGF*β*R axis activation and Smad singling, and another that depends upon CXCL12/CXCR4-axis activation, EGFR transactivation, and downstream signaling through MEK/ERK and PI3K/Akt pathways - all of which converge in the nucleus to promote the expression of multiple collagen genes (45, 46).

We previously reported that TGF*β* and CXCL12 induced or repressed a transcriptional molecular signature that was 75% similar and 25% dissimilar (27). There is evidence that both TGF*β* and CXCL12 may be acting upon the same set of bHLH (basic Helix-Loop-Helix) E-box and Egr-1/Egr-2 TFs that bind to consensus sequences in the promoters of the *COL1A1* and *COL1A2* and other genes (47, 48). To test this hypothesis we utilized the DGE profiles from the RNA-Seq experiments for both TGF*β* and CXCL12 treatment (27) and performed a CIE analysis using Ternary scoring statistic and merged tissue corrected ChIP-network.

The algorithm predicts many active regulators. Notably, **AHR, BHLHE40, TCF4, TCF12, ARNT, ARNTL, MYC**, and **NEUROG2** bHLH transcription factors, and **Egr-1, Egr-2** transcription factors are predicted by the algorithm as top putative regulators. We also examined the promoter of COL1A1 and COL1A2 and identified multiple binding sites for several of these TFs, including AHR, Egr-1, BHLHE40, ARNT, TCF4. In particular, Egr-1 has multiple binding sites in the promoters of these genes. Taken together, these results support the hypothesis that Egr-bHHL-family TFs can drive the expression of collagen genes in response to TGF*β* and CXCL12.

## CONCLUSION AND DISCUSSION

In this work, we present Causal Inference Engine (CIE), an integrated platform for identification and interpretation of active regulators of transcriptional response. The platform offers visualization tools and pathway enrichment analysis to map predicted regulators to Reactome pathways. We provide a parallelized `R`-package for fast and flexible directional enrichment analysis that can run the inference on any user provided custom regulatory network. Multiple inference algorithms are provided within the CIE platform along with regulatory networks from curated sources TRRUST and TRED as well as a causal protein-gene interactions derived from the STRINGdb. Importantly we provide a high confidence annotated causal transcriptional regulatory network by combining publicly available ChIP-Seq data with tissue specific gene expression data. Using a novel regularized gaussian graphical model, we softly enforce the TF-gene interaction identified by ChIP-Seq experiments in estimating the precision and partial correlation matrices form tissue gene-expression data, from which we drive tissue-specific annotated transcriptional regulatory networks. Further by merging the networks, we obtained a set of consistent TF-gene interactions that are universally applicable independent of the context. Benchmarks against the gold standard TRRUST database demonstrate that our approach is well able to recover mode of regulation with high accuracy. We demonstrated the utility of our approach in discovering known and novel biology using controlled *in vitro* over-expression studies as well as stem cell differentiation. Moreover, we demonstrated how our platform can be utilized to investigate specific biological hypotheses of transcriptional regulatory mechanisms in the context of fibroblast phenotypic plasticity in response to signaling events. Our approach and platform can be adopted for other settings, such as identifying candidate co-activators of specific transcription factors and reconstructing regulatory networks from single cell gene expression data. We hope that this platform provides the scientific community an open source alternative tool to interpret differential gene expression and to generate new biological insights. In the future, we plan to integrate additional networks and pathway inference methods in our platform. We also plan to pursue biological validation of our results on fibroblast phenotypic plasticity using ChIP-Seq methodologies.

## DATA AVAILABILITY

CIE is an open source collaborative initiative available in the GitHub repository: https://github.com/umbibio/CIE-R-Package All of the used databased can be downloaded through CIE web server via Download menu: https://markov.math.umb.edu/app/cie

## SUPPLEMENTARY DATA

Supplementary Data are available at NAR online.

## ACKNOWLEDGEMENTS

Authors would like to acknowledge Yasaman Rezvani for help with R-code.

