## Supplementary material for "Causal Inference Engine: A platform for directional gene set enrichment analysis and inference of active transcriptional regulators": SI

### Supplementary materials

| GTEx tissue | # Samples |
|---|---|
| bladder | 11 |
| breast | 218 |
| cervix | 11 |
| uterus | 90 |
| colon | 173 |
| liver | 136 |
| salivary gland | 70 |
| esophagus_mucosa | 285 |
| esophagus_gas | 193 |
| esophagus_muscle | 310 |
| prostate | 119 |
| stomach | 204 |
| thyroid | 355 |
| lung | 374 |
| kidney | 36 |

| | |
|---|---|
| Total | 2,585 |

**Supplemental Table 1.** Number of paired-end RNA-seq samples analyzed in each tissue using GTEx database

| Tissue | R-squared | Coefficient | #non-zero interactions in precision matrix |
|---|---|---|---|
| bladder | 0.479 | 2.8 | 196,643 |
| breast | 0.5448 | 2.6 | 199,317 |
| cervix | 0.5 | 2.6 | 192,844 |
| colon | 0.588 | 1 | 426,832 |
| esophagus_gas | 0.556 | 2 | 310,173 |
| esophagus_mucosa | 0.558 | 2.3 | 233,562 |
| esophagus_muscle | 0.563 | 2.2 | 250,310 |
| kidney | 0.513 | 2.9 | 230,009 |
| liver | 0.551 | 2.6 | 240,073 |
| lung | 0.568 | 2.7 | 194,549 |
| prostate | 0.543 | 2.4 | 269,741 |
| salivary | 0.531 | 2.5 | 280,796 |
| stomach | 0.57 | 2.2 | 232,734 |
| thyroid | 0.575 | 1.4 | 399,374 |
| uterus | 0.553 | 2.8 | 247,195 |

**Supplemental Table 2.** This table shows number of non-zero interactions per tissue which where the results of the proposed approach. The best R-squared value and its corresponding coefficient are also reported for each tissue.

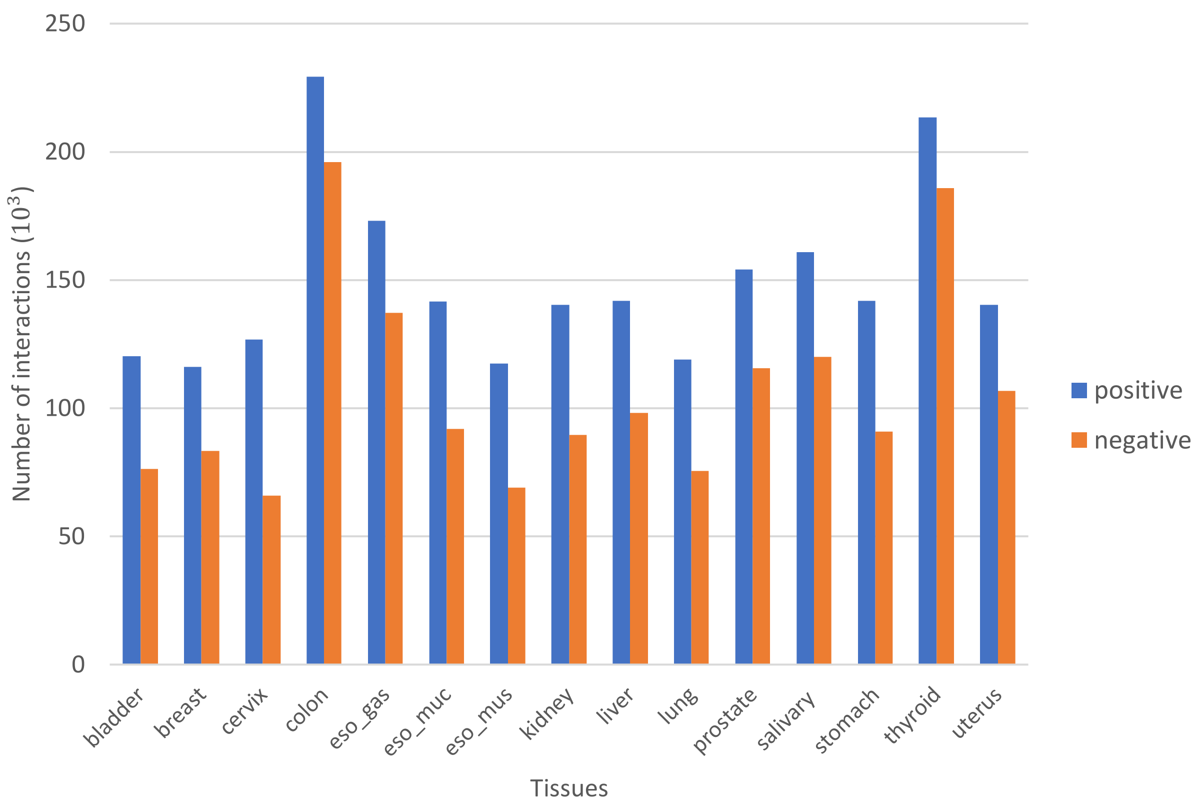


**Supplemental Figure 1.** Histogram of the interactions reported positive and negative signs across tissues

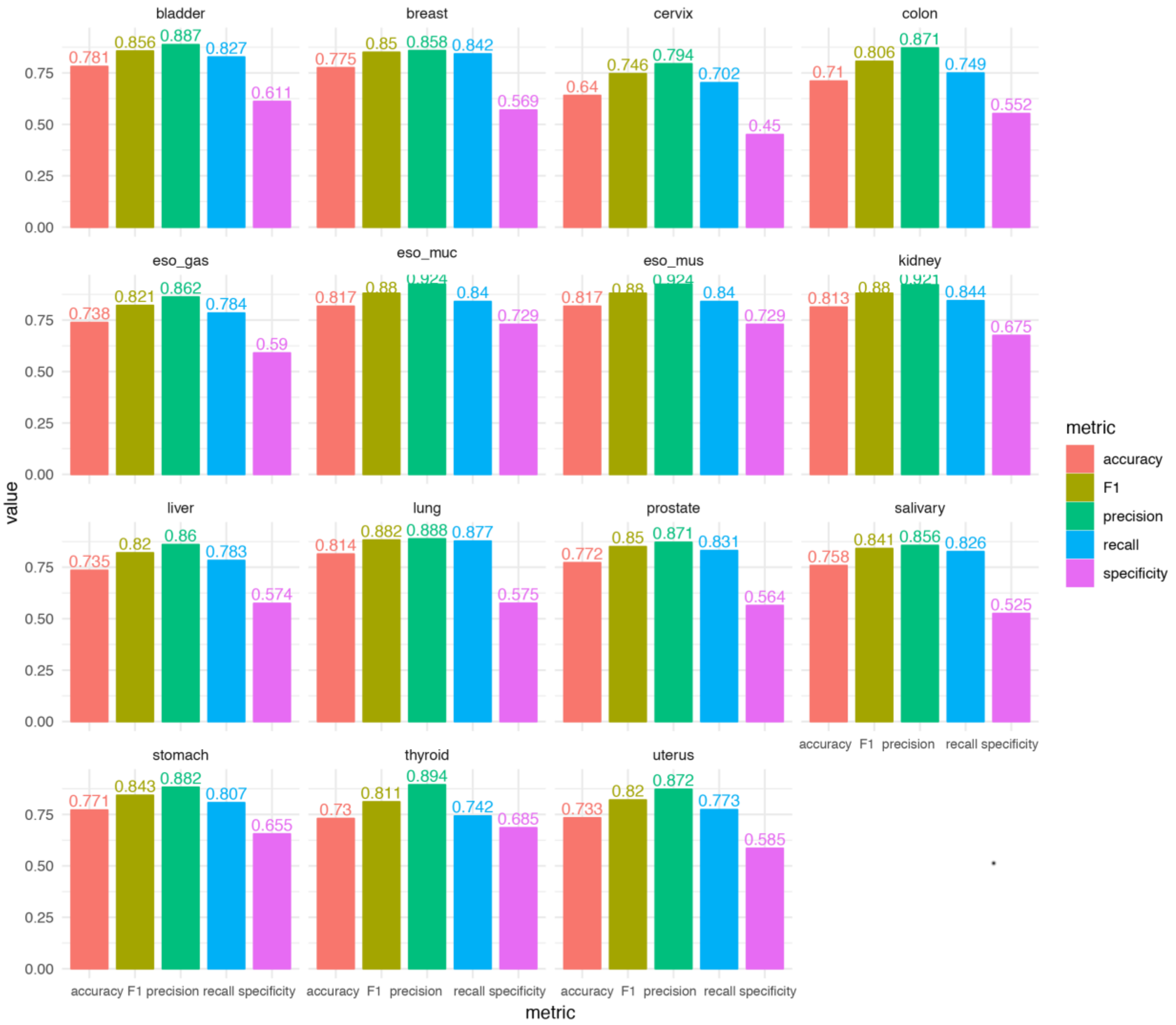


**Supplemental Figure 2.** Classification performance of our extracted signs compared to TRRUST reported mode of regulations across tissue
